# Regulation of Muscle Stem Cell Dynamics and Quiescence by Netrin-1 Cytoskeletal Signaling

**DOI:** 10.1101/2025.11.01.686042

**Authors:** Hsiao-Fan Lo, Yue Lu, David Sachs, Jean-Francois Cloutier, Frédéric Charron, Elizabeth Chen, Robert S. Krauss

## Abstract

The hallmark property of muscle stem cells (MuSCs) at homeostasis is quiescence. However, MuSCs have cellular protrusions that are heterogeneous, complex, and tipped with filopodia, all signs of motile structures. Such protrusions may serve as sensors of the MuSC niche. Through development of a novel ex vivo live imaging assay for MuSC protrusion dynamics, we report here the identification of a regulatory pathway in this process. The axon guidance cue Netrin-1 promotes MuSC protrusion outgrowth ex vivo in a manner dependent on its receptors Dcc and Neogenin, the small GTPase Rac1, and the actin-branching factor Arp2/3; this pathway is also required for Netrin-dependent axonal growth cone motility. Adult MuSC-specific genetic removal of Netrin-1 receptors, Rac1, and Arp2/3 each result in failure to maintain homeostatic protrusion lengths, spontaneous quiescence exit, and MuSC attrition in uninjured mice. These findings reveal an unanticipated level of morphological dynamism by a quiescent stem cell at homeostasis and link this phenomenon to preservation of the quiescent state.

---

Many adult mammalian tissues regenerate following injury, a property ascribable to tissue-specific stem cells^1, 2^. Prior to injury, such stem cells often reside in a state of quiescence, a reversible exit from the cell cycle that can persist for prolonged periods of time^3, 4^. Quiescence is actively promoted by signals from the stem cell niche and required for maintenance of the stem cell pool. However, how such stem cells sense and respond to a niche environment that is heterogeneous and dynamic is not well understood. Cells generally adopt morphologies that are adapted to such task^5^, but the in vivo morphology of most quiescent stem cell types is unknown.

Skeletal muscle stem cells (MuSCs, also called satellite cells) are a prime example of quiescent stem cell. Upon tissue injury, MuSCs are activated, proliferate, and differentiate to form new myofibers. They also self-renew, returning to quiescence and restoring the stem cell compartment. MuSC quiescence is an actively maintained state, dependent on multiple signals from the stem cell microenvironment, or niche^6–8^. MuSCs reside in direct contact with an associated myofiber and underneath an enwrapping basal lamina; this defines the immediate niche^9, 10^. Cells present outside the basal lamina, including fibroadipogenic progenitors (FAPs) and vasculature-associated cells, also serve as niche components^9, 10^. Niche factors required for MuSC quiescence can be divided into functional categories, including adhesion molecules (e.g., cadherins), morphogenetic factors (e.g., Notch pathway ligands), and GPCR ligands (e.g., neurotensin)^9, 11–18^.

MuSCs have traditionally been described as small, fusiform cells with a high nucleus-to-cytoplasm ratio^19, 20^. However, recent work has demonstrated that, in vivo, quiescent MuSCs have multiple, long cellular protrusions^21–24^. These structures are heterogeneous and complex, with distal subbranches and multiple filopodia^22, 25^. MuSCs with long protrusions display features of deeper quiescence than those with short or no protrusions^22, 23^. In response to muscle injury the protrusions rapidly retract, suggesting that they serve as sensors of the niche. MuSC protrusions have a core of microtubules surrounded by a sleeve of cortical F-actin, with distal localization of the actin-branching factor Arp2/3^22^. This morphology and cytoskeletal arrangement resembles that of neuronal axons and growth cones during CNS development^26, 27^. Similar to pathfinding axons, the Rac and Rho small GTPases are key regulators of MuSC protrusion dynamics, with high Rac1 activity promoting outgrowth and quiescence, and high levels of RhoA-ROCK signaling driving retraction and promoting subsequent MuSC activation events^22, 28^.

The morphology and cytoskeletal architecture of MuSC protrusions suggest that they might be dynamic, motile structures, even during quiescence, thereby providing MuSCs with the ability to survey their niche environment (we note that MuSC cell bodies are stationary at quiescence^21,29^). We have hypothesized that MuSC protrusion dynamics during quiescence are promoted by niche-derived signals acting via Rho family GTPases to maintain the range of protrusion lengths and morphologies observed at homeostasis, with some cues promoting outgrowth while other cues promote a level of retraction consistent with sustained quiescence^25^. Such factors are unknown, and new approaches are required for their identification.

Netrins are a small family of extracellular proteins related to laminins^30^. Netrin-1 promotes axon outgrowth and chemoattraction during CNS development^30, 31^. It does so via two paralogous receptors, Dcc and Neogenin (Neo1)^30, 32^. Neo1 also serves as a chemorepellent receptor for RGM family ligands and, with RGMs, as a coreceptor for BMP signaling^33^. In contrast, Dcc does not bind RGM and lacks these latter functions^34^. The intracellular regions of Dcc and Neo1 bind to a direct activator of Rac1 (the guanine nucleotide exchange factor, Arhgef7) and to the WAVE regulatory complex (WRC), a key regulator of actin cytoskeleton dynamics^35^. Rac1 activates WRC, which in turn signals via Arp2/3. The Netrin-1 → Dcc/Neo1 → Rac1 →WAVE → Arp2/3 pathway is required for netrin-mediated axon guidance in an evolutionarily conserved manner^36–38^.

We report here development of an ex vivo live imaging assay for MuSC protrusion dynamics and its use in identification of Netrin-1-dependent cytoskeletal signaling as a regulator of this process. Netrin-1 stimulates MuSC protrusion outgrowth ex vivo in a manner dependent on Dcc, Neo1, Rac1, and Arp2/3. Strikingly, MuSC-specific genetic removal of Dcc, Neo1, Rac1, and Arp2/3 each leads to a failure of MuSCs to maintain homeostatic protrusion lengths and to a break in MuSC quiescence in the absence of injury. Netrin-1–mediated regulation of MuSC protrusion dynamics is therefore linked to maintenance of quiescence. Furthermore, guidance cues, acting via changes to cell morphology, represent a new class of niche factor MuSC quiescence regulator.

## Results

### Development of an ex vivo assay for MuSC protrusion dynamics

To identify factors that regulate MuSC protrusion dynamics we developed an ex vivo assay involving short-term culture of muscle bundles (Fig. 1a and see Methods). Adult *Pax7^CreERT2^;R26^LSL-tdTomato^* (*Pax7^tdT^*) mice were used to label all MuSCs via Tamoxifen-induced expression of tdTomato. Extensor digitorum longus (EDL) muscles were excised and subjected to limited digestion with collagenase in DMEM. Simply excising a muscle is sufficient to initiate MuSC activation and, similar to the widely used single myofiber preparation^22^, this protocol activates MuSCs. To circumvent this problem, the medium was supplemented with the ROCK inhibitor, Y-27632 (Y27), to inhibit MuSC activation and preserve their protrusions.^22^ Myofiber bundles were then gently triturated from the EDL, washed to remove Y27, and imbedded in a type I collagen gel. These preparations were then subjected to live imaging by confocal microscopy. In these preparations, ∼85% of MuSCs fully retracted their protrusions and acquired a rounded appearance in ∼3 hours, a time frame similar to what is seen in vivo after acute injury^22^ (Fig. 1b, 1d, and Supplementary Video 1). We also observed MuSCs that had nearly completed protrusion retraction and then re-extending such structures outward (Fig. 1c and Supplementary Video 2). Approximately ∼15% of MuSCs extended protrusions, despite their being in an environment that promotes MuSC activation (Fig. 1d), suggesting that adult skeletal muscle harbors factors that promote MuSC protrusion outgrowth.

**Fig. 1.**
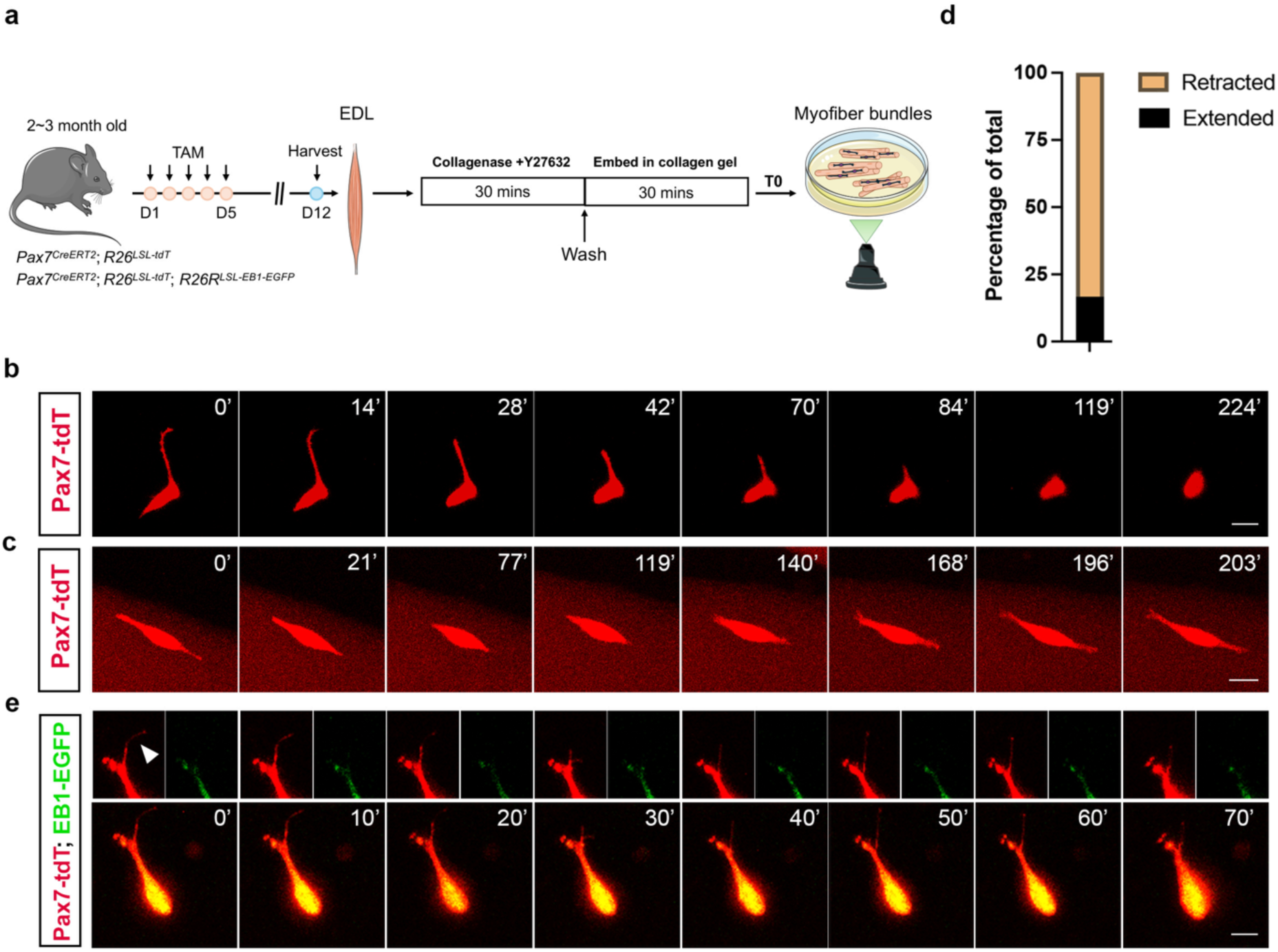
Live imaging of MuSC protrusion dynamics ex vivo. **a,** Schematic of the muscle bundle preparation protocol. **b,** Still images from live imaging of a tdTomato-labelled MuSC, within a muscle bundle, retracting its protrusion. Images were taken over a 4-hour time course. Scale bar, 10μm. **c,** Still images from live imaging of a tdTomato-labelled MuSC, within a muscle bundle, retracting and then extending protrusions. Images were taken over a 4-hour time course. Scale bar, 10μm. **d,** Percentage of MuSCs that retracted vs. extended protrusions. N=36 MuSCs from 11 mice. **e,** Still images from live imaging of a MuSC expressing both tdTomato and EB1-EGFP. Top panel shows individual tdTomato and EB1-EGFP signals, the bottom panel shows both signals and the entire cell. The arrowhead denotes a dynamic tdTomato^+^, EB1-EGFP^-^ filopodium. Scale bar, 10μm.

This assay also revealed dynamic behavior of filopodia at MuSC protrusion tips. We generated *Pax7^tdT^* mice that harbored a *R26^LSL-EB^*^1^*^-EGFP^* allele, allowing Tamoxifen-inducible expression of the microtubule plus-end binding protein EB1 fused to EGFP. Filopodia are F-actin-based, finger-like projections that are largely devoid of MTs. Filopodial structures labeled with tdTomato but not with EB1-EGFP were found to retract and extend at ends of MuSC protrusions (Fig. 1e and Supplementary Video 3). Elongated MuSCs that were neither retracting nor extending a full protrusion were still dynamic, extending filopodia repeatedly during a three-hour time course (Supplementary Video 4).

### Multiple muscle-resident cells express Netrin-1 and MuSCs express Netrin receptors

The morphological and cytoskeletal resemblance between MuSC protrusions and pathfinding axons raised the possibility that axon guidance cues are regulators of MuSC protrusion dynamics. We therefore analyzed existing single cell RNAseq datasets of muscle-resident mononuclear cells, as well as single nucleus RNAseq datasets of myonuclei, for expression of axon guidance factors and their receptors^39, 40^. We focus here on *Ntn1* (encoding Netrin-1) and the genes encoding the receptors responsible for its outgrowth and chemoattractant activities, *Dcc* and *Neo1*. *Ntn1* was expressed by FAPs, muscle-resident tenocytes, and endothelial cells (Extended Data Fig. 1a). Expression by myofibers was minimal^40^. To assess *Ntn1* expression in situ, we first used a mouse line with a *lacZ* reporter construct inserted into the *Ntn1* locus (Fig. 2a)^41, 42^. β-gal activity from this allele is produced as a fusion protein with N-terminal domains of Netrin-1; the protein is trapped in the secretory pathway leading to intracellular β-gal activity that faithfully reproduces the *Ntn1* expression pattern. Strong β-gal signal was observed in an elongated pattern radiating from the ankle myotendinous junction (MTJ), likely derived from intramuscular tenocytes. The MTJ at the knee also displayed intense β-gal activity. Strong β-gal signal was observed near the surface of the muscle from what are probably endothelial cells of larger blood vessels. Finally, there was more irregularly patterned staining throughout the EDL, likely derived from FAPs and/or capillary endothelial cells. All EDL muscles examined (n=5) showed a similar pattern of *Ntn1* expression, as did tibialis anterior (TA) muscles (Extended Data Fig. 1b). To further validate Netrin-1 protein production by these cells we performed immunofluorescence (IF) analyses on TA muscle sections with antibodies to Netrin-1 and markers of FAPs, tenocytes, and endothelial cells. Netrin-1 signal was associated with 29% of FAPs, identified with a *Pdgfra*-driven EGFP reporter; with 60% of tenocytes, identified with a *Scx*-driven EGFP reporter; and with endothelial cells, identified with antibodies to CD31 (Fig. 2b). It was difficult to fully accurately discern individual CD31^+^ endothelial cell boundaries, but the great majority of them were also positive for Netrin-1.

**Fig. 2.**
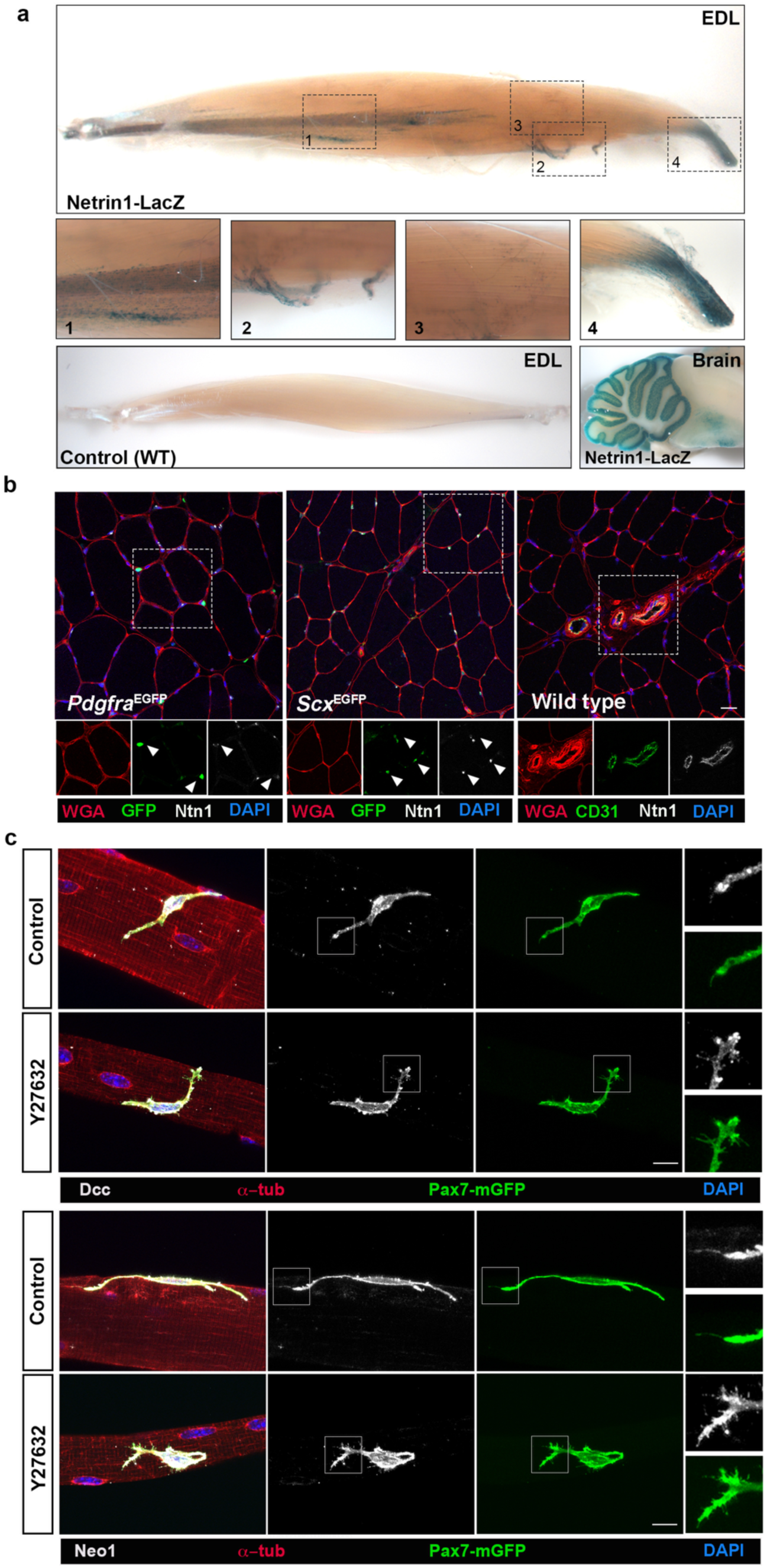
Expression of Netrin-1 by MuSC niche cells and Netrin-1 receptors by MuSCS. **a,** In situ analysis of β-gal activity on an EDL muscle from a *Ntn1-lacZ* reporter mouse. Boxed areas 1-4 are enlarged to reveal different patterns of staining. Based on scRNAseq analyses (Extended Data Fig. 1), the likely cell types expressing β-gal activity are: Box 1, muscle-resident tenocytes radiating from the ankle myotendinous junction; Box 2, endothelial cells from a surface blood vessel; Box 3, FAPs and capillary endothelial cells; and 4, tenocytes from the knee myotendinous junction. An EDL from a wild-type (WT) mouse stained for β-gal activity is shown as a negative control, a section through the brain of an adult mouse, with strong β-gal in the cerebellum, is shown as a positive control. EDL muscles from 5 independent mice yielded similar β-gal expression patterns. **b,** Immunofluorescence analysis of Netrin-1 (Ntn1) expression in sections of TA muscles. Wheat germ agglutinin (WGA) outlines myofibers, a *Pdgfra^EGFP^*reporter identifies FAPs, a *Scx^EGFP^* reporter identifies tenocytes, and antibody to CD31 identifies endothelial cells. Boxed images highlight Ntn1 protein signal associated with the given cell type marker. Scale bar, 20μm. **c,** Immunofluorescence analysis of Netrin-1 receptor expression in MuSCs. *Pax7^mG^* mice were generated as in Figure 1a. Single myofibers from *Pax7^mG^* mice were prepared in the absence (Control) or presence of Y27632, which preserves protrusion length and protrusion tip structures, including filopodia. Myofibers were immunostained with antibodies to α-tubulin (α-tub) to reveal myofibers, to mGFP to reveal MuSC membranes, and Dcc or Neo1 as indicated. Boxed areas are shown at higher magnification at far right. Note that in the presence of Y27632, Dcc and Neo1 localization is enriched at tips of MuSC protrusions. Scale bars, 10μm.

We next assessed expression of *Dcc* and *Neo1* by different cell types in skeletal muscle, particularly MuSCs. scRNAseq datasets indicated that *Neo1* is broadly expressed, including by FAPs, various vasculature-associated cells, and MuSCs (Extended Data Fig. 1a); some myonuclei also expressed *Neo1* transcripts^40^. In contrast, *Dcc* expression was barely detected (Extended Data Fig. 1a). This latter result is consistent with extremely low levels of *Dcc* mRNA in bulk RNAseq analyses of MuSCs^43, 44^. To assess protein production, IF analysis was used on single myofibers, an ex vivo preparation that maintains MuSCs in their immediate niche. We recently modified this technique such that MuSC protrusions, which usually retract during myofiber isolation, were partially preserved.^22^ Single EDL myofibers were prepared under control conditions and in the presence of Y27, which further preserves in vivo protrusion length and complex cellular structures at protrusion tips, including filopodia.^22^ To enhance visualization of such structures, these experiments used *Pax7^CreERT^*^2^*;R26^mTmG^* (*Pax7^mG^*) mice, labelling MuSCs with membrane-targeted (myristoylated) GFP (mGFP). MuSCs expressed Neo1 and, despite its extremely low mRNA levels, Dcc proteins (Fig. 2c; we note here that the Dcc protein signal is specific, data shown below). In Y27-treated preparations, these receptors were enriched at protrusion tips, similar to what is observed in filopodia-bearing axonal growth cones.^45^ Taken together, these results indicate that Netrin-1 is produced by multiple cell types in adult skeletal muscle, and MuSCs produce the chemoattractant Netrin-1 receptors, Dcc and Neo1.

### Netrin-1 stimulates MuSC protrusion outgrowth ex vivo

To test whether MuSCs respond to Netrin-1 with outgrowth of protrusions, we exploited the muscle bundle live imaging assay. Recombinant Netrin-1 was added uniformly to the collagen gel in which muscle bundles are imbedded, and MuSC protrusion dynamics were monitored by confocal microscopy over 4 hours. Netrin-1 strongly and dose-dependently increased the percentage of MuSCs that extended, rather than retracted, their protrusions (Fig. 3a, 3b, and Supplementary Video 5). An evolutionarily conserved pathway required for Netrin’s outgrowth and attractant function in axon guidance requires the small GTPase, Rac1, and the actin-branching factor, Arp2/3.^36, 38^ Inclusion in the collagen gel of small molecule inhibitors of Rac (NSC23766) or Arp2/3 (CK-666) each inhibited Netrin-1–stimulated MuSC protrusion outgrowth (Fig. 3a, 3b, and Supplementary Videos 6 and 7). MuSCs in these ex vivo muscle bundles begin the activation process during the preparation, but longer MuSC protrusions correlate with a more quiescent phenotype.^22, 23^ Accordingly, we assessed expression of Fos, a marker of early MuSC activation.^22, 46–48^ Approximately 60% of MuSCs in control bundle preparations were Fos^+^ after 4 hours (Fig. 3c, 3d). This percentage was reduced to <40% by treatment with recombinant Netrin-1, in a manner dependent on Rac and Arp2/3 activities (Figure 3c, 3d). Therefore, Netrin-1 treatment promoted extension of MuSC protrusions and diminished MuSC activation, and both activities relied on signaling via the cytoskeleton.

**Fig. 3.**
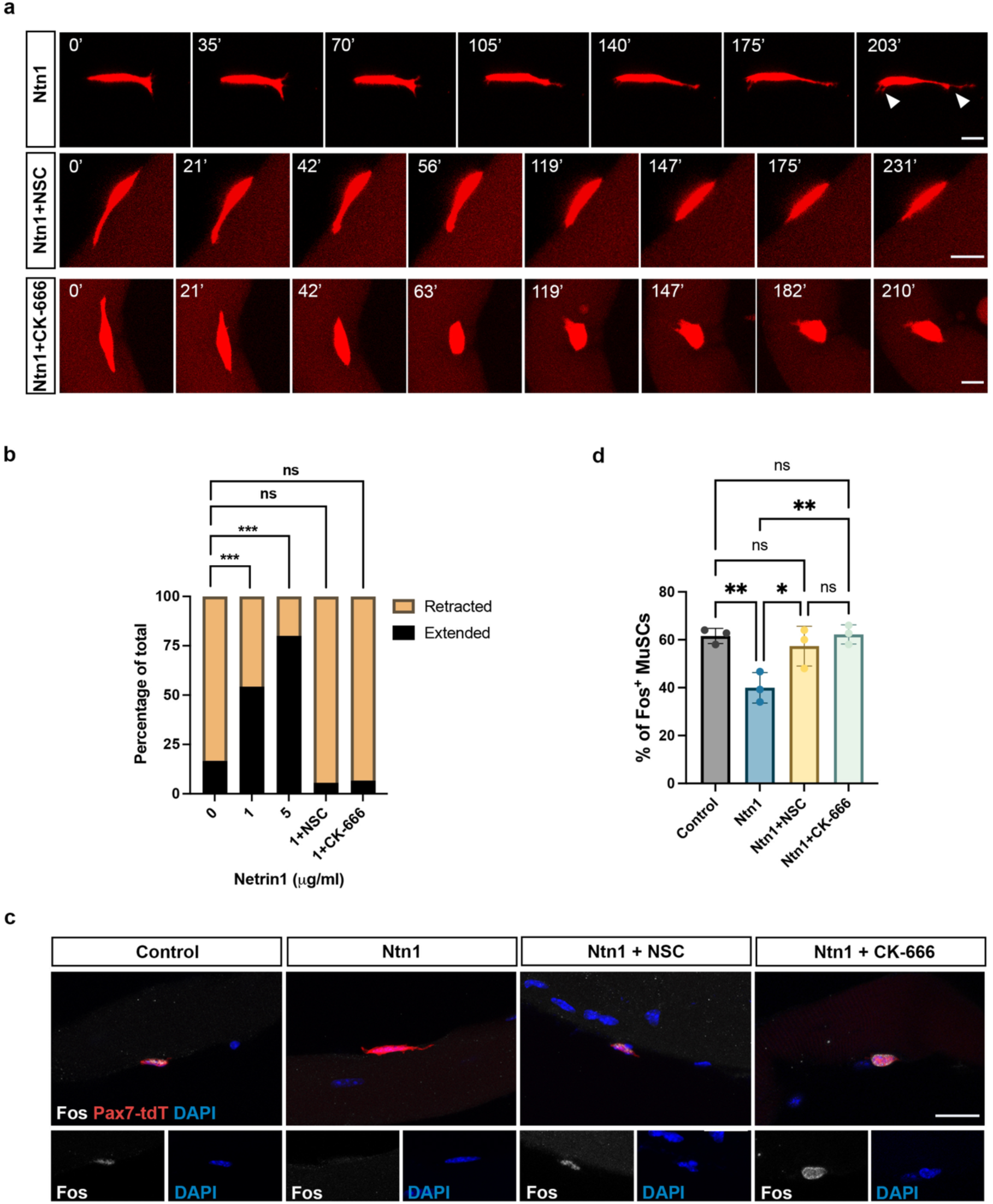
Netrin-1 stimulates MuSC protrusion outgrowth ex vivo. **a,** Mice were generated as in Figure 1a. Top panel, still images from live imaging of a tdTomato-labelled MuSC, within a muscle bundle treated with recombinant Netrin-1 (Ntn1), extending a protrusion Arrowheads indicate sites of protrusion extension. Middle panel, still images from live imaging of a tdTomato-labelled MuSC, within a muscle bundle treated with recombinant Ntn1 and the Rac inhibitor NSC, retracting a protrusion. Bottom panel, still images from live imaging of a tdTomato-labelled MuSC, within a muscle bundle treated with recombinant Ntn1 and the Arp2/3 inhibitor CK-666, retracting a protrusion. Scale bars, 10μm. **b,** Percentage of MuSCs that retracted vs. extended protrusions under the indicated conditions. For 1μg/ml Ntn1, N=35 MuSCs from 6 mice; for 5μg/ml Ntn1, N=15 MuSCs from 4 mice; 1μg/ml Ntn1 plus NSC inhibitor (50mM), N=18 MuSCs from 3 mice; 1μg/ml Ntn1 plus Arp2/3 inhibitor (200 mM), N=15 MuSCs from 4 mice. **c,** MuSCs within bundles incubated under the indicated conditions immunostained for tdTomato and Fos. Scale bar, 20μm. **d,** Percentage of Fos^+^ MuSCs under the indicated conditions. ≥50 MuSCs were quantified per EDL, N=3 mice for each treatment.

### *Dcc* and *Neo1* are required for MuSC quiescence and maintenance

To provide genetic evidence for regulation of MuSC dynamics and quiescence in vivo by Netrin-1 signaling, we genetically removed Dcc and Neo1 specifically from MuSCs. This approach is preferable to using *Ntn1* mutants because *Ntn1* and *Neo1* are both expressed by multiple MuSC niche cell types. This is similar to the hematopoietic stem cell (HSC) niche, wherein multiple niche cell types produce Netrin-1, and it regulates interactions both between niche cells and HSCs and also between niche cells themselves.^49, 50^ It is possible that mutation of *Ntn1* in various niche cells could disrupt signaling within MuSC niche compartments and thereby affect MuSCs indirectly by altering niche properties. In contrast, mutation of *Dcc* and *Neo1* specifically in MuSCs permits analysis of stem cell-autonomous effects of these receptors.

Mice carrying homozygous floxed alleles for *Dcc* or *Neo1*, or both genes together, were crossed with *Pax7^tdT^* and/or *Pax7^mG^*mice, yielding *Dcc* cKO, *Neo1* cKO, and *Dcc;Neo1* double cKO (dcKO) mice, respectively (Fig. 4a). Effects of loss of either receptor were initially determined on single EDL myofiber preparations analyzed by IF for Dcc or Neo1 and the appropriate fluorescent reporter. *Dcc* cKO MuSCs displayed loss of Dcc protein expression (Fig. 4b), confirming that the IF signal was specific even though *Dcc* mRNA levels in RNAseq datasets are very low. Furthermore, the average protrusion length was reduced by >70% (Fig. 4d). Very similar results were observed with *Neo1* cKO and *Dcc;Neo1* dcKO MuSCs (Fig. 4c and 4d). *Dcc* cKO MuSCs still expressed Neo1 protein and vice versa (Extended Data Fig. 2a). Because MuSC protrusions begin retracting during preparation of single myofibers, we also studied the effects of loss of these receptors on protrusion length in vivo via tissue clearing of EDL muscles. *Dcc* cKO, *Neo1* cKO, and *Dcc;Neo1* dcKO MuSC protrusion lengths were reduced by 46%, 43%, and 53% from that in control MuSCs, respectively; none of the three mutant lines were statistically different from each other (Fig. 4e and 4f). TA muscles from control and the three mutant lines were also examined using immediate in situ fixation of the excised muscle; the effects on protrusion length were very similar to those seen with cleared EDL muscles (Extended Data Fig. 2b, 2c).

**Fig. 4.**
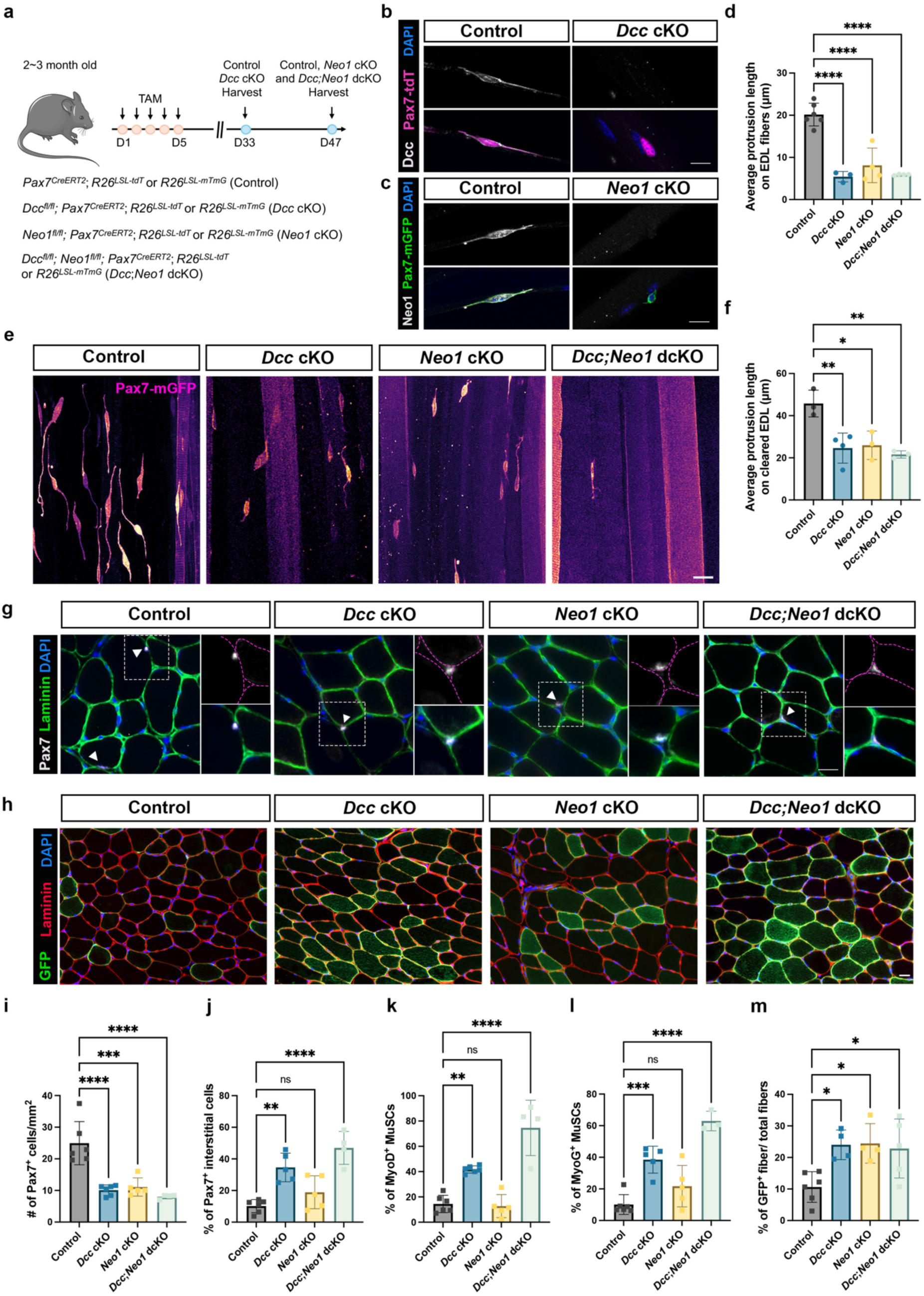
Dcc and Neo1 are required for MuSC quiescence and maintenance. **a,** Scheme for generation and analysis of *Dcc* cKO, *Neo1* cKO, and *Dcc;Neo1* dcKO mice. *Dcc* cKO mice were harvested 4 weeks after Tamoxifen (TAM) treatment and *Neo1* cKO and *Dcc;Neo1* dcKO mice were harvested 6 weeks after TAM treatment. The additional two weeks were used for the latter because Neo1 protein perdured for that long after TAM (see Methods). **b,** Immunofluorescence analysis of Dcc expression and protrusion length in MuSCs from control and *Dcc* cKO mice. Single EDL myofibers from *Pax7^tdT^* mice were prepared and immunostained for Dcc. Note the absence of Dcc signal in the *Dcc* cKO MuSC. Scale bars, 10μm. **c,** Immunofluorescence analysis of Neo1 expression and protrusion length in MuSCs from control and *Neo1* cKO mice. Single EDL myofibers from *Pax7^mG^* mice were prepared and immunostained for Neo1. Note the absence of Neo1 signal in the *Neo1* cKO MuSC. Scale bars, 10μm. **d,** Quantification of average MuSC protrusion length on single EDL myofibers from control, *Dcc* cKO, *Neo1* cKO, and *Dcc;Neo1* dcKO mice. Each point represents the average protrusion length of at least 50 MuSCs from an individual mouse. **e,** Images of cleared EDL muscles from *Dcc* cKO, *Neo1* cKO, and *Dcc;Neo1* dcKO mice. Scale bar, 20μm. **f,** Quantification of average MuSC protrusion length in cleared EDL muscles from control, *Dcc* cKO, *Neo1* cKO, and *Dcc;Neo1* dcKO mice. Each point represents the average protrusion length of at least 50 MuSCs from an individual mouse. **g,** Immunofluorescence analysis of TA muscle sections from control, *Dcc* cKO, *Neo1* cKO, and *Dcc;Neo1* dcKO mice. Sections were stained with antibodies to Pax7 and laminin, which outlines myofibers. Myofiber outlines are highlighted with dashed magenta lines in the insets to clearly reveal whether MuSCs are under the basal lamina (Control and *Neo1* cKO) or in the interstitium (*Dcc* cKO and *Dcc;Neo1* dcKO). Scale bars, 20μm. **h,** Immunofluorescence analysis of TA muscle sections from control, *Dcc* cKO, *Neo1* cKO, and *Dcc;Neo1* dcKO *Pax7^mG^* mice. Myofibers positive for expression of mGFP indicate fusion of MuSCs has occurred. Scale bars, 20μm. **i-l,** Quantification of numbers of Pax7^+^ cells, interstitial Pax7^+^ cells, MyoD^+^ cells, and MyoG^+^ cells in sections represented in (**f**) and Extended Data Fig. 3. **m,** Quantification of numbers of mGFP^+^ myofibers observed in sections represented in panel (**g**).

Genetic removal of factors required for maintenance of MuSC quiescence leads to activation in the absence of injury, often followed by cell attrition.^14, 51–53^ Lack of, or short, protrusions are associated with MuSC activation^22, 23^, and we therefore quantified Pax7^+^ MuSCs on sections of TA muscle from all three mutant lines. The number of Pax7^+^ cells was reduced in *Dcc* cKO, *Neo1* cKO, and *Dcc;Neo1* dcKO mice by 59%, 55%, and 68% from control values, respectively; similar to the changes seen with protrusion lengths, the degree of MuSC loss was not statistically different between the three mutant lines (Fig. 4g and 4i). The percentage of Pax7^+^ cells present in the interstitium (i.e., outside the basal lamina, having exited the immediate niche) was significantly elevated in *Dcc* cKO and *Dcc;Neo1* dcKO, but not *Neo1* cKO, mutants (Fig. 4g, 4j). We next quantified cells expressing the lineage determinant and MuSC activation marker, MyoD, and the differentiation marker, myogenin (MyoG). MyoD^+^ and MyoG^+^ cells were elevated, relative to Pax7^+^ cells, in *Dcc* cKO and *Dcc;Neo1* dcKO mice but this ratio in *Neo1* cKO mice was not different from controls (Fig. 4k, 4l, and Extended Data Fig. 3a, 3b).

To identify underlying mechanisms of MuSC depletion in Netrin-1 receptor mutants, we analyzed whether they underwent either premature differentiation and fusion with myofibers or cell death via apoptosis. To address the former, we used lineage tracing of MuSCs from *Pax7^mG^* control, *Dcc* cKO, *Neo1* cKO, and *Dcc;Neo1* dcKO mice by scoring the percentage of mGFP^+^ myofibers in sections of TA muscles.^16^ Each mutant line displayed a >2-fold increase in mGFP^+^ myofibers, consistent with the conclusion that some MuSCs from these mice were depleted by fusion with myofibers (Fig. 4h and 4m). To test whether any mutant MuSCs underwent apoptosis, TA sections were stained with antibodies to Pax7 and cleaved Caspase-3 (cCasp3). The percentage of Pax7^+^ MuSCs that were also cCasp3^+^ was elevated in each mutant line, though only dcKO mice rose to a level of p<0.05 by one-way ANOVA (Extended Data Fig. 3c, 3d). We noticed, however, that the vast majority of cCasp3^+^/Pax7^+^ MuSCs were located in the interstitial space, irrespective of genotype (80-95% of such cells, across genotypes). These results suggest that loss of Netrin-1 receptor-mutant MuSCs by apoptosis is likely to be quantitatively less important than loss by premature differentiation and fusion. Nevertheless, elevated levels of interstitial MuSCs are found in *Dcc* cKO and *Dcc;Neo1* dcKO mutants, and MuSCs that escape the niche are more prone to depletion by apoptosis. MuSC attrition in Netrin-1 receptor-mutant MuSCs therefore occurs via multiple mechanisms.

In summary, genetic removal of Dcc and Neo1 each led to short MuSC protrusions and attrition of MuSCs, but this was not exacerbated when the two receptors were removed simultaneously. Loss of Dcc, but not Neo1, led to exit of MuSCs from the niche and expression of MyoD and MyoG. However, simultaneous loss of Dcc and Neo1 did not exacerbate these phenotypes beyond what was seen with loss of Dcc alone. Taken together, Dcc and Neo1 are both required for homeostatic maintenance of quiescent MuSCs, but they do not display an obvious additive or synergistic effect.

### Dcc and Neo1 are each required for Netrin-1–stimulated MuSC protrusion outgrowth

Dcc and Neo1 are both Netrin-1 receptors involved in axon outgrowth and chemoattraction^30, 32^, but Neo1 is also a receptor for RGM family ligands and as a coreceptor for BMP ligands, functions Dcc lacks.^33, 34^ To address the roles of Dcc and Neo1 in Netrin-1 signaling in MuSCs, we performed live imaging on muscle bundle preparations from *Dcc* and *Neo1* cKO mice. Control MuSCs responded to recombinant Netrin-1 by extending cellular protrusions in a dose-dependent manner. In contrast, neither *Dcc* nor *Neo1* cKO MuSCs displayed protrusion outgrowth when treated with Netrin-1 (Fig. 5a – 5c and Supplemental Videos 8 and 9). Therefore, both Dcc and Neo1 receptors are required for Netrin-1’s ability to promote MuSC protrusion outgrowth.

**Fig. 5.**
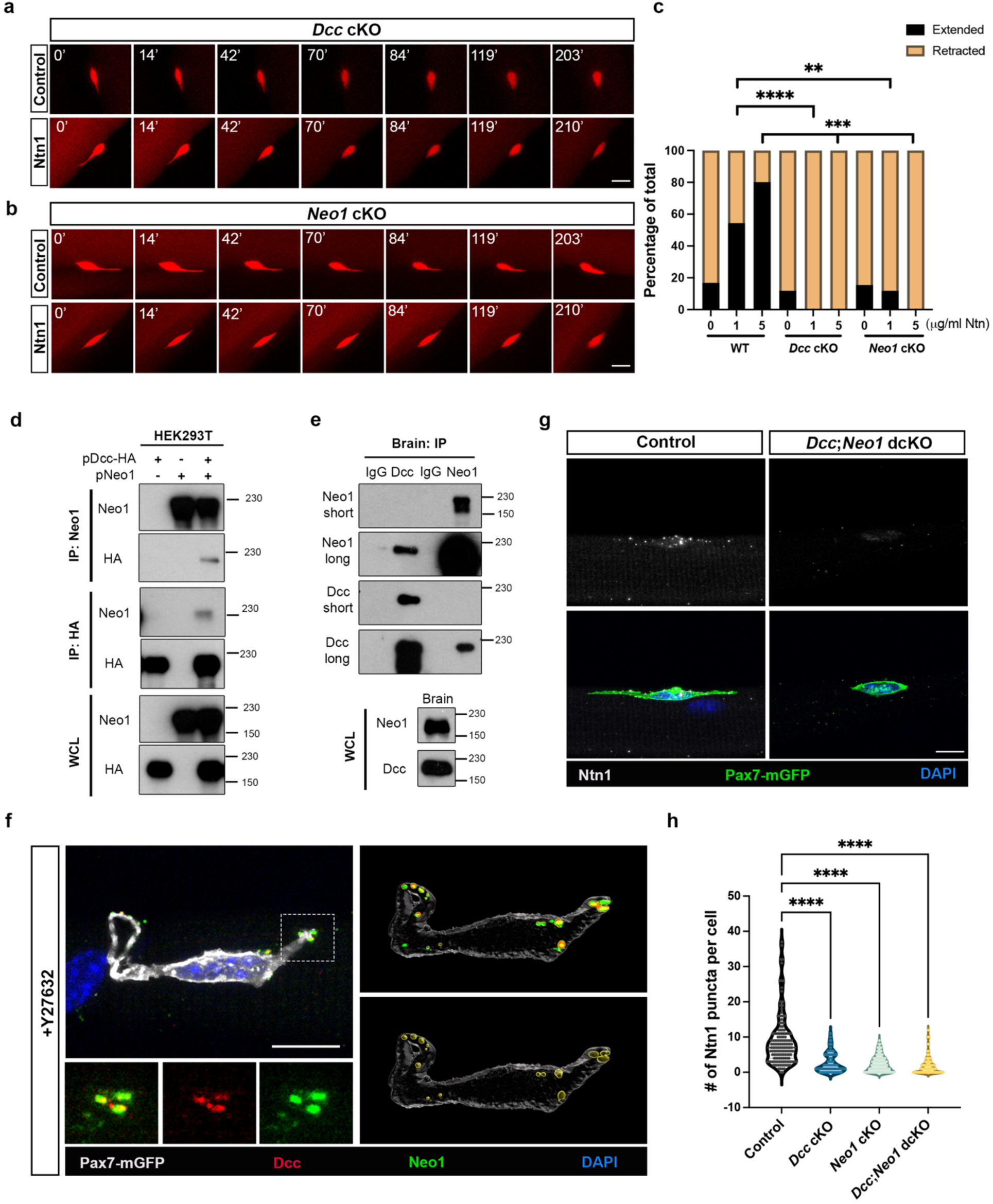
Dcc and Neo1 are required for Netrin-1 – stimulated MuSC protrusion outgrowth and associate with each other. **a,** Mice were generated as in Figure 4a. Still images from live imaging of a tdTomato-labelled, *Dcc* cKO MuSC, within a muscle bundle treated with either vehicle (Control) or recombinant Netrin-1 (Ntn1). Note that the Ntn1-treated MuSC does not extend a protrusion. Scale bar, 10μm. **b,** Still images from live imaging of a tdTomato-labelled, *Neo1* cKO MuSC, within a muscle bundle treated with either vehicle (Control) or recombinant Ntn1. Note that the Ntn1-treated MuSC does not extend a protrusion. Scale bar, 10μm. **c,** Percentage of WT, *Dcc* cKO, and *Neo1* cKO MuSCs that retracted vs. extended protrusions under the indicated conditions. N=17 MuSCs from 3 *Dcc* cKO mice (0μg/ml Ntn1); N=14 MuSCs from 3 *Dcc* cKO mice (1μg/ml Ntn1); N=13 MuSCs from 3 *Neo1* cKO mice (0μg/ml Ntn1); N=17 MuSCs from 3 *Neo1* cKO mice (1μg/ml Ntn1). **d,** HEK293T cells were transiently transfected with the HA-tagged Dcc and/or Neo1 expression vectors, and cell lysates were immunoprecipitated with HA or Neo1 antibodies, then immunoblotted with the indicated antibodies. Whole cell lysates (WCL) were blotted as controls. **e,** Whole brain lysates from P14 mice were immunoprecipitated with antibodies to Dcc, Neo1, or IgG isotype control and immunoblotted with the indicated antibodies. Short and long refer to length of exposure time. Whole brain cell lysates (WCL) were blotted as controls. **f,** Confocal immunofluorescent analysis of Dcc and Neo1 localization in MuSCs. Single myofibers from *Pax7^mG^* mice were prepared in the presence of Y27632. Myofibers were immunostained with antibodies to mGFP, Dcc, and Neo1. The boxed area is shown at higher magnification below and reveals overlapping localization of Dcc and Neo1 in puncta at the tip of a MuSC protrusion. The right-hand panels display analyses of the image with Imaris software. In the lower images, the areas circled in yellow are regions of Dcc-Neo1 colocalization, Scale bar, 10μm. **g,** Immunofluorescence analysis of Ntn1 puncta associate with MuSCs on single myofibers from control and *Dcc;Neo1* dcKO mice. Scale bar, 10μm. **h,** Quantification of Ntn1 puncta on MuSCs on single myofibers from control, *Dcc* cKO, *Neo1* cKO, and *Dcc;Neo1* dcKO mice

Guidance of commissural axons during spinal cord development is the archetypal system for Netrin-1 activity.^30, 54^ *Dcc* and *Neo1* both function in commissural axon guidance, but only *Dcc* mutants display a loss of function phenotype (a function for *Neo1* is revealed by its mutation in addition to mutation of *Dcc*, and the two together account for all Netrin-1-dependent outgrowth and chemoattractant activity in this system).^32^ Structural studies show that Dcc and Neo1 form Netrin-bound homomeric receptor complexes.^32, 55^ This is consistent with the genetic analyses of commissural axon guidance that suggest they function in the same process, but independently of one another.^32, 55^ Our observation that individual mutation of either *Dcc* or *Neo1* resulted in: 1) shortened MuSC protrusions in vivo; and 2) loss of Netrin-1-responsiveness ex vivo, demonstrated a need for both *Dcc* and *Neo1* in these cells. Furthermore, the phenotype of *Dcc;Neo1* dcKO mice was not significantly more severe than the single cKO mice. This scenario raised the possibility that Dcc and Neo1 form heteromeric receptor complexes. To address this possibility, we co-transfected vectors encoding Neo1 and epitope-tagged Dcc into HEK293T cells and performed reciprocal co-immunoprecipitations on cell lysates. Antibodies to Neo1 brought down Dcc and antibodies to the Dcc epitope tag brought down Neo1 (Fig. 5d). To assess whether this interaction occurs endogenously, extracts of P14 mouse brains were immunoprecipitated with antibodies to Dcc or Neo1 and immunoblotted for the other. Dcc and Neo1 again co-precipitated with one another (Fig. 5e). Finally, we performed IF for both proteins on single EDL myofiber preparations. Confocal microscopy revealed that, in MuSCs, Dcc and Neo1 co-localized in puncta (Fig. 5f). Colocalization of Dcc and Neo1 was also demonstrated with Imaris software analysis of confocal images (Fig. 5f).

During guidance of specific neuronal axons during *C. elegans* development, UNC-6 (Netrin) forms puncta on their growth cones in an UNC-40 (Dcc)-dependent manner.^56^ Accordingly, we assessed whether Netrin-1 formed puncta on MuSCs and what the receptor requirements were. IF analysis of single myofibers revealed Netrin-1 puncta on control MuSCs and these were strongly diminished in *Dcc* cKO, *Neo1* cKO, and *Dcc;Neo* dcKO MuSCS (Fig. 5g, 5h). As *Ntn1* is not appreciably expressed by myofibers or MuSCs themselves, these results demonstrate that niche-derived Netrin-1 reaches, and is present on, MuSCs, dependent on the presence of both Dcc and Neo1. The results in Fig. 5, combined with the lack of additive or synergistic effects in *Dcc;Neo* dcKO MuSCs (Fig. 4), are consistent with the notion that Dcc and Neo1 act together in Netrin-1 signaling in MuSCs, perhaps as heteromeric Netrin-1 receptor complexes.

### Arp2/3 is required for maintenance of MuSC protrusions and maintenance

Arp2/3 promotes branched actin formation and is present at the tips of growing axons and MuSC protrusions.^22, 26, 27^ It is also important for Netrin-1–stimulated axon guidance and outgrowth of MuSC protrusions (^30^ and Fig. 3). To assess whether Arp2/3 activity is required in homeostatic MuSCs, we genetically removed from these cells *Arpc2* (Fig. 6a), which encodes an essential subunit of the Arp2/3 complex. Single EDL myofibers were isolated from control and *Arpc2* cKO mice and immunostained for CD34 to measure MuSC protrusion length. The average protrusion length of *Arpc2* cKO MuSCs was reduced by 70% (Fig. 6b, 6c).

**Fig. 6.**
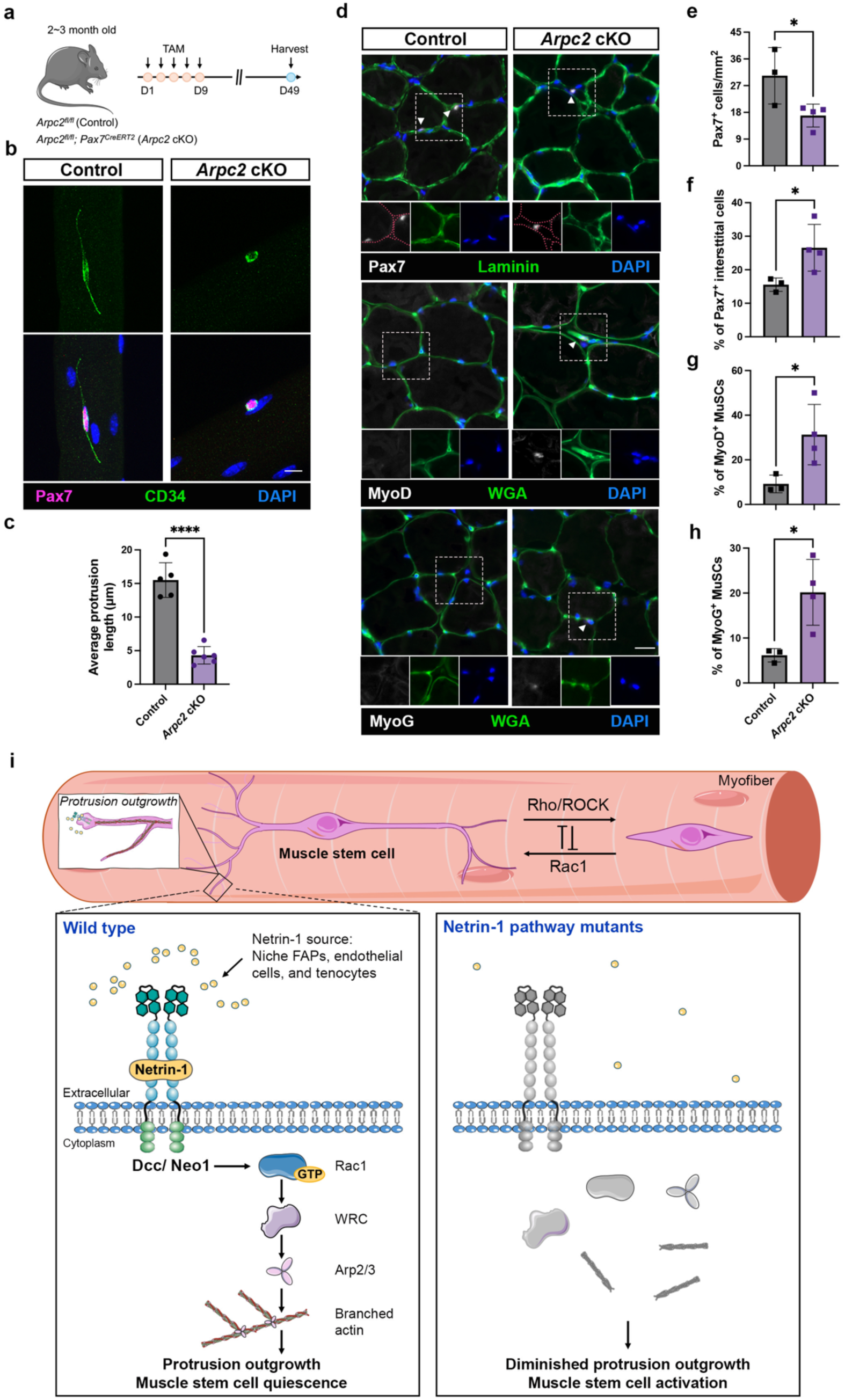
Loss of Arp2/3 results in short MuSC protrusions and MuSC attrition. **a,** Scheme for generation and analysis of *Arpc2* cKO mice. **b,** Immunofluorescence analysis of protrusion length in MuSCs from control and *Arpc2* cKO mice. Single EDL myofibers were prepared and immunostained for Pax7 to identify MuSCs and CD34 to measure protrusions. Scale bar, 10μm. **c,** Quantification of average MuSC protrusion length on single EDL myofibers from control and *Arpc2* cKO mice. Each point represents the average protrusion length of at least 50 MuSCs from an individual mouse. **d,** Immunofluorescence analysis of TA muscle sections from control and *Arpc2* cKO mice. Sections were stained with antibodies to Pax7, MyoD, or MyoG, and myofibers were outlined by either staining with antibodies to laminin or with WGA. Scale bar, 20μm. **e-h,** Quantification of numbers of Pax7^+^ cells, interstitial Pax7^+^ cells, MyoD^+^ cells, and MyoG^+^ cells in sections represented in (**c**). **i,** Model of quiescent MuSC protrusion dynamics. In this model, MuSCs are morphologically mutable, ranging from cells with long, complex protrusions to those with fusiform shape. The balance is predicted to be governed by the relative levels of Rac1 vs. RhoA/ROCK signaling regulated by niche cues.^24^ One such cue is Netrin-1 which signals via Rac1 to promote protrusion outgrowth and quiescence. MuSCs lacking Netrin-1 signaling factors fail to maintain homeostatic protrusion lengths and break quiescence in the absence of injury. FAPs, fibroadipogenic progenitors. See text for further details.

Furthermore, the number of Pax7^+^ cells in TA sections was reduced by 56% (Fig. 6d, 6e). In analyses of TA sections, loss of ArpC2 resulted in elevated percentages of interstitial Pax7^+^ cells, MyoD^+^ cells, and MyoG^+^ cells (Fig. 6d, 6f-h). These numbers are very similar to those seen with *Dcc* mutants (Fig. 4). These results indicate that MuSCs have an ongoing requirement for Arp2/3-dependent production of branched actin to maintain quiescence.

## Discussion

The MuSC niche provides multiple extracellular cues that are independently required for MuSC quiescence and, in turn, maintenance of the stem cell population.^9^ These can be placed into categories based on their mechanism of action; e.g., adhesion molecules and morphogenetic factors. Quiescent MuSCs possess elaborate cellular protrusions that share morphological and cytoskeletal features with pathfinding axons during CNS development.^22, 23^ Here we report that the axon guidance cue Netrin-1 signals via cytoskeletal regulators to stimulate MuSC protrusion outgrowth, and that these factors are required in mice for both homeostatic maintenance of protrusions and quiescence. Netrin-1 represents the first member of a new category of quiescence-promoting MuSC niche factor: guidance cues. Furthermore, these findings reveal an unanticipated level of morphological dynamism by a quiescent stem cell at homeostasis and link this phenomenon to preservation of the quiescent state.

In adult muscles at homeostasis, MuSC protrusions are highly heterogeneous.^22, 23^ Our findings provide genetic evidence that these cellular structures are motile during quiescence. The results are best explained by a model whereby niche cues signal to quiescent MuSCs to maintain a dynamic state of protrusion length and morphology, some cues promoting outgrowth, while others promote a degree of retraction consistent with maintenance of quiescence.^25^

Netrin-1 signaling drives protrusion outgrowth, thereby helping to maintain this equilibrium. Loss of Netrin-1 signaling shifts the range to shorter protrusion lengths, often to the point where, presumably, MuSCs aberrantly enter the activation process (Fig. 6i). An alternative possibility is that MuSC protrusion morphologies in vivo are unique but fixed, with Netrin-1 signaling required to maintain this state. However, Netrin-1 is sufficient to drive MuSC protrusion outgrowth in muscle bundles ex vivo, and this activity is dependent on downstream signaling factors (Rac1 and Arp2/3) that are also required for Netrin-1’s ability to promote axon outgrowth and guidance. These same factors are required for maintenance of homeostatic MuSC protrusion lengths and quiescence. Taken together, the results strongly favor the former explanation.

Regulation of quiescent MuSC protrusions holds similarity to mechanisms whereby pathfinding axons are guided during development. Netrin-1, the major chemoattractant cue in the developing CNS, signals via multiple downstream pathways.^30^ One well-established pathway, in which Netrin-1 signals via Dcc to Rac1 and WRC to promote Arp2/3 activity, is required in vivo for attraction of axons to the midline in an evolutionarily conserved manner, including in *Drosophila*, mice, and humans.^36, 38, 57, 58^ We find that Netrin-1’s ability to promote MuSC protrusion outgrowth ex vivo is dependent on Dcc and its paralog Neo1, Rac1, and Arp2/3. Furthermore, genetic removal of *Dcc*, *Neo1*, *Rac1*, or *Arpc2* (a component of the Arp2/3 complex) from MuSCs in vivo each lead to short protrusions and loss of quiescence (^22^ and this study). These observations reveal that an ancient developmental signaling pathway that promotes patterning of the CNS also controls dynamic behavior of adult stem cells. Mutations in humans that compromise this pathway produce axon pathfinding defects that result in corpus collosum agenesis and congenital mirror movement disorder (CMM).^59^ Humans with mutations in *CDH2* (encoding N-cadherin) show an overlapping clinical spectrum, including these axon pathfinding defects.^60^ Like Netrin-1, N-cadherin functions as an axon guidance factor in *Drosophila* and mice.^61–65^ We have previously reported that N-cadherin localizes to MuSC protrusion tips, and MuSC-specific mutation of *Cdh2* in mice results in short protrusions and loss of quiescence.^13, 22^ Therefore, multiple extracellular cues that are required for specific axon guidance events also regulate MuSC protrusion dynamics and quiescence. Together, these findings reveal that growth cone-bearing axons and quiescent MuSC protrusions have significant morphological and mechanistic similarities.

How quiescent stem cells sense their niche is not well understood. Quiescent MuSC protrusions rapidly retract after muscle injury, leading to the hypothesis that they serve as cellular sensors of the MuSC niche.^22, 23, 25^ While this seems likely, and it is in keeping with other injury-sensitive cell types in which cellular protrusions play such a role (e.g., microglia^66^), one may ask what benefits motility brings to a MuSC niche sensor function. Dynamic MuSC protrusions could provide a surveillance role, allowing MuSCs to probe the surface of myofibers. The surface of myofibers is likely to be non-uniform and changeable with muscle contraction.

Dynamic protrusions might allow MuSCs to sense a larger area of a myofiber surface that is itself dynamic. Changes to the MuSC cytoskeleton occur extremely rapidly after an activation stimulus^22^, and we have suggested that the first signals after injury that MuSCs detect are changes to the biomechanical properties of the niche.^25^ Consistent with this notion, MuSC-specific genetic removal of the mechanosensitive ion channel TRPM7 results in delays to Rho activation, protrusion retraction, and MuSC activation.^67^ Dynamic MuSC protrusions may provide an efficient means for these stem cells to distinguish injury-induced changes to the niche from the normal dynamics of the niche, such as changes that occur due to myofiber contraction. Inability of MuSCs to maintain an appropriate degree of morphological dynamism, as evidenced here through loss of Netrin-1 signaling, leads to a break in quiescence. This finding argues that MuSC protrusion motility is closely linked to cytoskeletal structures that are in turn sensitive to quiescence-maintaining niche cues.

Many other adult stem cell types exist in a state of niche-dependent quiescence, and our findings with MuSCs are likely to be relevant to some of these. For example, Netrin-1 is expressed by multiple HSC niche cell types, and it signals via Neo1 to promote HSC quiescence in vivo.^50^ Treatment of HSCs in vitro with recombinant Netrin-1 for 48 hr promoted quiescence-associated properties but the signaling pathways directly downstream of Neo1 are not known. Although HSCs have been studied in great detail, their in vivo morphology is not well established, nor are the in vivo morphologies of many types of quiescent adult stem cell. HSCs do, however, display limited motility within their niche, and they are sensitive to cytoskeletal regulation and capable of extending cellular protrusions when treated in vitro with other niche factors.^68–71^ Our findings provide a plausible mechanistic basis for Netrin-1/Neo1 signaling in HSC quiescence. The concept that quiescent, adult stem cells can be morphologically mutable offers flexible mechanisms for such cells to dynamically probe their niche environment. This in turn can provide active maintenance of quiescence and rapid cellular responses upon tissue injury.

## Acknowledgements

We thank Artur Kania for providing Dcc conditional mutant sperm, Zhuhao Wu for providing *Ntn1*-lacZ reporter mice and advice and help with tissue clearing, Arun Narasimhan for help with Imaris software analysis, Camelia Gonzalez and Angelina Kramer for technical assistance, and Gab Kardon and Paul Wassarman for critically reading the manuscript. This work was funded by National Institute of Arthritis and Musculoskeletal and Skin Diseases grants to R.S.K. (R01AR070231 and R21AR08463) and to E.H.C. (R01AR085116), by a National Institute of General Medical Sciences grant to E.H.C. (R35GM136316), and by a Canadian Institutes of Health Research grant to J.F.C. Microscopy was performed at the Microscopy and Advanced Imaging CoRE at the Icahn School of Medicine at Mount Sinai (P30CA196521 from the Tisch Cancer Institute at the Icahn School of Medicine at Mount Sinai). For Figure 1a, the image was adapted from Servier Medical Art (https://smart.servier.com/), licensed under CC BY 4.0 (https://creativecommons.org/licenses/by/4.0/).

## Author Contributions

Conceptualization, H.-F.L., F.C., and R.S.K.; Methodology, H.-F.L. and R.S.K.; Reagents, J.F.C.; Validation and Formal Analysis, H.-F.L. and Y.L.; Investigation, H.-F.L. and Y.L.; Writing – Original Draft, R.S.K.; Writing – Review & Editing, all authors; Visualization, H.-F.L.; Supervision, R.S.K. and E.H.C.; Funding Acquisition, R.S.K. and E.H.C.

## Competing Interests

The authors declare no competing or financial interests.

## Methods

### Animals

Mice were housed and maintained in accordance with recommendations set in the Guide for the Care and Use of Laboratory Animals of the National Institutes of Health. All animal protocols were approved by the Icahn School of Medicine at Mount Sinai Institutional Animal Care and Use Committee (IACUC). Male and female mice were used for all experiments. Mice were genotyped by PCR using toe genomic DNA.

Isolated sperm from *Dcc^f/f^* mice^72^ and *Neo1^f/f^* mice^73^ were used for in vitro fertilization of oocytes isolated from C57BL/6J mice (JAX). IVF was performed by the Mount Sinai Mouse Genetics Shared Research Facility. *Pax7^CreERT2^* ^74^, *R26^LSL-tdT^* ^75^, and *R26^LSL-mTmG^*^76^ mice were from JAX. *Scx^EGFP^*mice^77^ were provided by Dirk Hubmacher, *Pdgfra^H2B-EGFP^* mice^78^ were provided by Phil Soriano, and *Ntn1^lacZ^* mice^42^ were provided by Zhuhao Wu. *Arpc2^f/f^* mice^79^ were used as previously described^80^. *R26R^LSL-EB1-EGFP^* mice were from RIKEN Center for Biosystems Dynamics Research.

Adult mice (2-3 months of age) were injected intraperitoneally for five consecutive days with 200 μL of a 12.5mg/mL tamoxifen solution (Toronto Research Chemicals, T006000) dissolved in corn oil (Thermo Scientific Chemicals, 405435000). Control (*Pax7^CreERT2^;R26^LSL-tdT^*and *Pax7^CreERT2^;R26^LSL-mTmG^*) mice were harvested one week after tamoxifen delivery. Neo1 protein displayed long perdurance, taking more than 4 weeks post-tamoxifen delivery to become undetectable by immunostaining of MuSCs on single fibers. *Neo1* cKO mice and *Dcc;Neo1* dcKO mice were therefore harvested six weeks after tamoxifen administration. All other mice were harvested four weeks after tamoxifen delivery. To estimate the recombination efficiencies *Dcc* and *Neo1* cKO mice, we made the assumption that any MuSCs that were no longer detectable via Pax7 staining had undergone full recombination, i.e. lacking expression of Dcc or Neo1 protein and degradation of perduring protein via homeostatic turnover. Minimum recombination efficiency by this definition is 59% for *Dcc* cKO mice and 55% for *Neo1* cKO mice, the percentage of MuSCs lost in these lines. Of the 41% of remaining *Dcc* cKO MuSCs, only 21.6% were Dcc immunoreactive; of the 45% of remaining *Neo1* cKO MuSCs, only 13.1% were Neo1 immunoreactive. These observations yielded an estimated 91.2% recombination efficiency in *Dcc* cKO mice and 94.1% recombination efficiency in *Neo1* cKO mice.

### Single myofiber isolation and immunofluorescence analysis

Single myofibers were isolated from EDL muscles as previously described.^22^ Isolated EDLs were incubated in DMEM media (supplemented with 10% horse serum, 10mM HEPES, and 1% Penicillin/Streptomycin) containing type I collagenase (2.8 mg/mL; Gibco 17100-017) in a 37°C shaking water bath for 53-55 minutes, followed by gentle trituration with a wide-mouth glass pipette. After trituration, plates were placed in a 37°C incubator for 10-15 minutes. For fixation, 25-30 straight myofibers were transferred from the trituration plate to a FACS tube. Myofibers were washed with PBS for 1 min, fixed with 4% paraformaldehyde for 10 minutes, and then washed with PBS three times for 5 minutes each. Myofibers intended for immunofluorescence experiments were then incubated in 0.2% Triton X-100 in PBS (PBST) for 10 minutes and blocked with 10% normal donkey serum for 1 hour at room temperature. Subsequently, primary antibodies were added and incubated overnight at 4°C. After washing with PBST, myofibers were incubated with secondary antibodies for one hour at room temperature. Nuclei were counterstained with DAPI, then mounted with Prolong Diamond antifade mountant (Thermo Fisher, P36970). Primary antibodies were: goat anti-Dcc (R&D Systems, AF-844), rabbit anti-Dcc (Abcam, ab273570), goat anti-Neogenin1 (R&D Systems, AF-1079), goat anti-Netrin1 (R&D Systems, AF-1109), chicken anti-GFP (AvesLab, GFP-1010), rabbit anti-α-tubulin (Abcam, ab18251), and mouse anti-Fos (Santa Cruz Biotechnology, sc-166940). Biotin-conjugated, anti-CD34 mouse monoclonal (RAM34; ThermoFisher, 13-0341-82) was detected with Alexa Fluor™ 488 conjugated streptravidin (Invitrogen, S11223).

In documenting interaction between Dcc and Neo1 in MuSCs associated with single myofibers, image analysis and colocalization quantification were performed using Imaris 11.0 software. Raw image files (lif. format) were imported into Imaris, using the File Converter software, for processing. To restrict the analysis to the region of interest, a Surface object was created to mask the relevant muscle stem cell area (mGFP was used for this analysis). The Surface object was created with the surface creation wizard using the following parameters: Enable Region Of Interest = false, Enable Region Growing = false, Enable Tracking = false, Enable Classify = false, Enable Shortest Distance = false, Enable SurfacesROI = false, Enable Smooth = true, Surface Grain Size = 0.180 µm, Enable Eliminate Background = false, Active Threshold = true, Manual Threshold Value = 7, Filter Surfaces with option “Number of Voxels Img=1” above 1.00e4 voxels. Next, masked channels were created using mGFP (Green Surface, pseudocolor is gray in Fig. 5f) as a mask on Dcc (Magenta channel, pseudocolor is red in Fig. 5f) and Neo1(Gray channels, pseudocolor is green in Fig. 5f): The mask intensity was set to the original channel’s value for inside the Green Surface, while it was set to 0 outside the Green Surface. Colocalization was performed on the Masked Magenta and Masked Gray channels using the following parameters: threshold Magenta = 77.000, threshold Gray = 95.000. The results from the colocalization analysis were as follows - number of colocalized voxels – 452, Pearson’s coefficient in dataset volume – 0.7171, original Manders’ coefficient A – 0.9203, original Manders’ coefficient B – 0.8263, threshholded Manders’ coefficient A – 0.0942, thresholded Manders’ coefficient B – 0.0525. A new Coloc Surface object was then created, using the surface creation wizard on the Coloc Channel, with the following parameters: Enable Region Of Interest = false, Enable Region Growing = false, Enable Tracking = false, Enable Classify = false, Enable Shortest Distance = false, Enable SurfacesROI = false, Enable Smooth = true, Surface Grain Size = 0.180 µm, Enable Eliminate Background = false, Manual Threshold Value = 25, Filter Surfaces with option “Number of Voxels Img=1” above 10.0 voxels. Finally, secondary masked channels were created using the Mask channels option with mask intensity set to the original channel value inside the Coloc Surface and, for outside the surface, for all three channels, Coloc channel, Masked Magenta (Dcc) and Masked Gray (Neo1) channels. These secondary masked channels were using in the representative figures.

### Live imaging of MuSCs in muscle bundles

Muscle bundles were isolated from the EDL muscle. The preparation was similar to isolating single myofibers, but the incubation time for type I collagenase digestion was reduced to 30 minutes. The ROCK inhibitor Y27632 (10μM; StemCell Technologies, 72304), was included in the media during digestion to preserve MuSC protrusions. After digestion, the EDL was gently triturated with a wide-mouth glass pipette to obtain 25-100 myofiber bundles. Muscle bundles were then transferred to a microscope slide and washed with an aqueous solution of 1.5 mg/ml rat tail collagen I (Thermo Fisher, A1048301), thereby removing the digestion media and Y27632. Then bundles were then gently transferred to a glass-bottom dish (Ibidi, 81218-200) containing 1.5 mg/ml rat tail collagen I solution. This mixture was allowed to solidify for 30 min at 37 °C, yielding a gel with embedded muscle bundles. For ex vivo treatments, recombinant Netrin-1 protein (1 μg/ml or 5 μg/ml final concentrations, dissolved in PBS; R&D systems, 1109-N1-025/CF) was included in the collagen I solution before adding it to the glass-bottom. The Rac1 inhibitor NSC23766 (50μM, dissolved in water; Sigma-Aldrich, SML0952) and the Arp2/3 complex inhibitor CK666 (200μM, dissolved in DMSO; Sigma-Aldrich, 182515) were used similarly. Vehicle control experiments displayed no effect on MuSC protrusion dynamics. Time-lapse imaging was performed using a Leica SP8 STED microscope equipped with a stage-top incubation chamber with environmental controls. Three to six fields of view were recorded under a 20X objective lens, with a time interval of 7 minutes and a Z-step size of 1-2 μm for 240 minutes. MuSC protrusion lengths were measured at the 182-minute time point and compared to the initial time of imaging (time zero).

### Tissue clearing

Mice were anesthetized by 3-3.5% isoflurane inhalation delivered via a precision vaporizer (VetEquip 911103). Cardiac perfusion was performed with a peristaltic pump (Gilson F155006) at 8-10 ml/minute, with room temperature (RT) PBS for 1 minute, followed by RT 4% PFA for 7 minutes, then RT PBS for 3 minutes. All buffers contained 10 μg/ml heparin during cardiac perfusion. EDL muscles wer dissected and post-fixed in 4% PFA at 4 °C overnight, then washed 3 times in PBS at RT for 1 hour. The fixed samples were delipidated following a modified Adipo-Clear protocol^81, 82^. muscles were washed in a methanol/B1N buffer gradient (0%, 20%, 40%, 60%, 80% methanol; B1N buffer is H_2_O/0.1% Triton X-100/0.3 M glycine, pH7) for 15 minutes at each step, twice with 100% methanol for 15 minutes, once with dichloridemethane for 30 minutes, and 3 times with 100% methanol for 15 minutes. Samples were rehydrated by washing in a reverse methanol/B1N gradient (80%, 60%, 40%, 20%) for 15 minutes at each step. This was followed by washing with B1N for 1 hour and with 5% DMSO/0.3M Glycine/PTxwH overnight. The next day, muscles were washed 3 times with PTxwH for 15 minutes (PTxwH is PBS/0.1% Triton X-100/0.05% Tween 20/ 2 mg/ml heparin). Muscles (2-3 EDL muscles per group) were immunostained with 1μg rabbit anti-GFP antibody (Rockland, 600-401-215) in 200μL PTxwH at RT for 2 days, washed 3 times in PTxwH at RT for 1 hours, then stained with 10μg Atto 647-conjugated goat anti-rabbit antibody (Rockland, 611-156-122) in PTxwH at RT for 2 days. Samples were then washed 3 times in PTxwH at RT for 1 hour and further fixed in 1% PFA at 4°C overnight, washed 3 times in PTxwH at RT for 1 hour, then blocked in B1N at RT overnight. The next day, samples were washed in PTxwH, then bleached in 0.3% H_2_O_2_ at 4°C overnight, washed 3 times in 20mM PB at RT for 1 hour and twice with 25% 2,2’ thiodiethanol/ 10mM PBS at RT, once for 2 hours and then overnight. Finally, the samples were equilibrated with ACB buffer (PB, 2,2’ thiodiethanol and Iohexol) with refractive index adjusted to 1.53 using 2,2’-thiodiethanol and mounted on concave microscope slides for imaging. Images were taken by Leica SP8 STED microscope.

### Isolation and analysis of in situ fixed TA muscles and immunohistochemistry of TA sections

TA muscles were isolated then fixed immediately with 4% PFA for 35 minutes then washed 3 times in PBS. Fixed TA muscles were placed in PBST (0.5% Triton X-100) to soften them and Moria forceps were used to tease muscle bundles from the tendon, followed by isolation of single myofibers from bundles under the dissecting microscope. Myofibers were then transferred to microscope slides and mounted with Prolong diamond antifade mountant. For immunohistochemistry on TA sections, the isolated TA muscle was immediately embedded in 10% Tragacanth gum (Thermo Scientific Chemical, AAA18502) and snap-frozen in liquid nitrogen-cooled 2-methylbutane/isopentane (Fisher Scientific, O3551-4). 10μm-thickness cryosections were processed with a Leica CM3050S cryostat. Cryosections were fixed in 4% PFA for 15 minutes at RT, then permeabilized with ice-cold methanol at −20 °C for 5 minutes. Antigen retrieval was performed on cryosections using a 0.01M sodium citrate buffer, pH 6.0. Cryosections were washed with PBST (0.2% TritonX-100), then incubated in blocking buffer (5% BSA in PBST) for 2 hours. Primary antibodies were applied to sections overnight at 4 °C. and included rabbit anti-Laminin (Sigma-Aldrich, L9393), mouse anti-Pax7 (Developmental Studies Hybridoma Bank, Pax7-c), mouse anti-Myogenin (Developmental Studies Hybridoma Bank, F5D), mouse anti-MyoD (BD Biosciences, ab554130), rabbit anti-cleaved caspase3 (Cell signaling, 9661), chicken anti-GFP (AvesLab, GFP-1010), and rabbit anti-CD31 (Abcam, ab124432). The next day, cryosections were washed with PBST and incubated with secondary antibodies, including Wheat Germ Agglutinin, Alexa Fluor 488 Conjugate (Invitrogen, W11261), Goat anti-mouse IgG1 cross-adsorbed secondary antibody, Alexa Fluor 647 (Thermo Fisher, A-21240), Goat anti-rabbit IgG (H+L) cross-adsorbed secondary antibody, Alexa Fluor 488 (Thermo Fisher, A-11008), Donkey anti-Rabbit IgG (H+L) Highly Cross-Adsorbed Secondary Antibody, Alexa Fluor™ 555 (Thermo Fisher, A-31572), at RT for one hour. After washing with PBS and counterstaining with DAPI, slides were mounted with Prolong Diamond antifade mountant then examined under a 20X objective lens on a Zeiss AxioImager Z2 microscope.

### Co-immunoprecipitation

HEK293T cells were transfected with pcDNA3-hDcc-3XHA and pEF-Cyto-Neogenin1-Myc for 48 hours (plasmids courtesy of Fred Charron). The cells were then harvested in lysis buffer containing 50 mM Tris/HCl (pH 8.0), 150 mM NaCl, 2 mM EDTA, 10% glycerol, 0.5% Nonidet P40, 1 mM DTT, phosphatase inhibitor cocktail, and protease inhibitor cocktail (Sigma–Aldrich) for immunoprecipitation and immunoblotting with anti-HA (Sigma-Aldrich, H3663) and anti-Neogenin1 (R&D Systems, AF-1079) antibodies. The interaction of endogenous Dcc and Neo1 was examined using brain tissue from P14 mice. Brain lysate was immunoprecipitated with anti-Dcc (R&D Systems, AF-844) and anti-Neo1 antibodies, then washed five times with lysis buffer. The immunoprecipitated proteins were analyzed by immunoblotting with anti-Dcc and anti-Neo1 antibodies.

### β-galactosidase (LacZ) staining of muscle

Brain, EDL and TA muscles were collected from adult *Ntn1^lacZ^*mice. Muscles were also collected from wild type mice as a negative control. Brains and muscles were fixed in 4% paraformaldehyde for 1 hour, then washed 3 times for 5 minutes with PBS. Samples were incubated overnight in staining solution (1mg/ml X-gal, 0.02% NP-40, 5mM K_3_FeCN, 5mM K_4_FeCN, and 2mM MgCl_2_ in PBS) at 37°C. The next day, samples were transferred to a new tube and washed 3 times with PBS. Stained samples were photographed with a Jenoptik ProgRes C3 camera attached to Nikon SMZ 1500 stereomicroscope.

### Analysis of RNA sequencing databases

Data were downloaded from scMuscle, a set of all mononuclear cells in skeletal muscle^38^, and from Myoatlas, a snRNAseq database of myonuclei^39^, and converted to Seurat objects. Immune cells were pre-filtered from the data, cells were clustered, and UMAP plots were constructed to display single cell expression of selected genes using the default clustering parameters set by the database authors.

### Image Quantification

Protrusion length quantification was performed with Fiji software as described previously.^22^ At least 50 independent MuSCs were quantified per mouse on EDL single myofibers, TA myofibers, and tissue clearing samples. For quantification of cell numbers on TA cryosections during homeostasis, 10 random fields were captured with a 20X lens from at least 3 mice per group. More than 50 GFP-positive cells were quantified from n=3 mice to identify the localization of Netrin-1 on TA muscles from *Pdgfra^EGFP^* and *Scx^EGFP^* mice. For quantification of Netrin-1 puncta, at least 30 MuSCs were measured from n≥3 mice of control, *Dcc* cKO, *Neo1* cKO, and dcKO EDL myofibers.

### Statistical analysis

For live imaging experiments, n≥3 mice were harvested for each treatment. Fig. 1d and Fig. 3b shared the same control group. The significance of percentages of extending and retracting MuSC protrusions was analyzed by Fisher exact test between each group. A two-tailed unpaired t-test with Mann-Whitney test was used for the number of Netrin-1 puncta. For all other experiments, data with more than two independent groups were analyzed by one-way ANOVA, multiple comparison between each group. Graph and P values were generated using GraphPad Prism and used to determine the level of varying p values between experimental groups. N.S. (not significant) = p > 0.05, * = p < 0.05, ** = p < 0.01, *** = p < 0.001, **** = p < 0.0001.

## Extended Data Figure Legends

**Extended Data Fig. 1.**
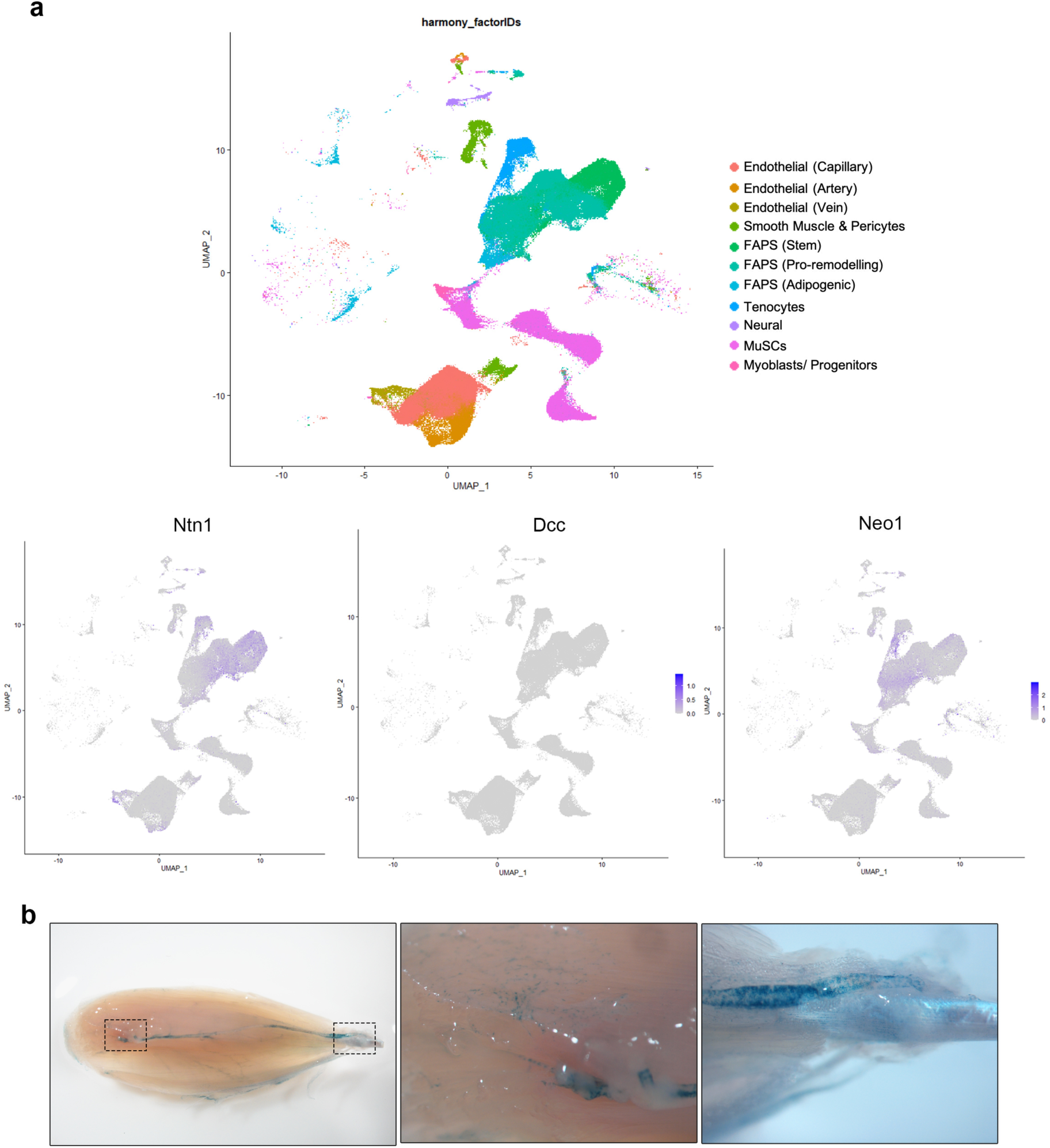
Expression of *Ntn1, Dcc*, and *Neo1* in scRNAseq datasets and *Ntn1* in adult muscle. **a,** Single cell RNA-sequencing analysis of resident mononuclear cells in adult mouse skeletal muscle^39^ revealed 11 different cell types after Harmony integration, and they are color-coded here (note: hematopoietic cells have been removed from the analysis). Expression of *Ntn1*, *Dcc*, and *Neo1* are documented. **b,** In situ analysis of β-gal activity on an TA muscle from a *Ntn1-lacZ* reporter mouse. Boxed areas are enlarged to reveal different patterns of staining. Based on scRNAseq analyses, the likely cell types expressing β-gal activity are: Left box, endothelial cells from a surface blood vessel, and FAPs and capillary endothelial cells distributed throughout the muscle; Right box, muscle-resident tenocytes radiating from the ankle myotendinous junction.

**Extended Data Fig. 2.**
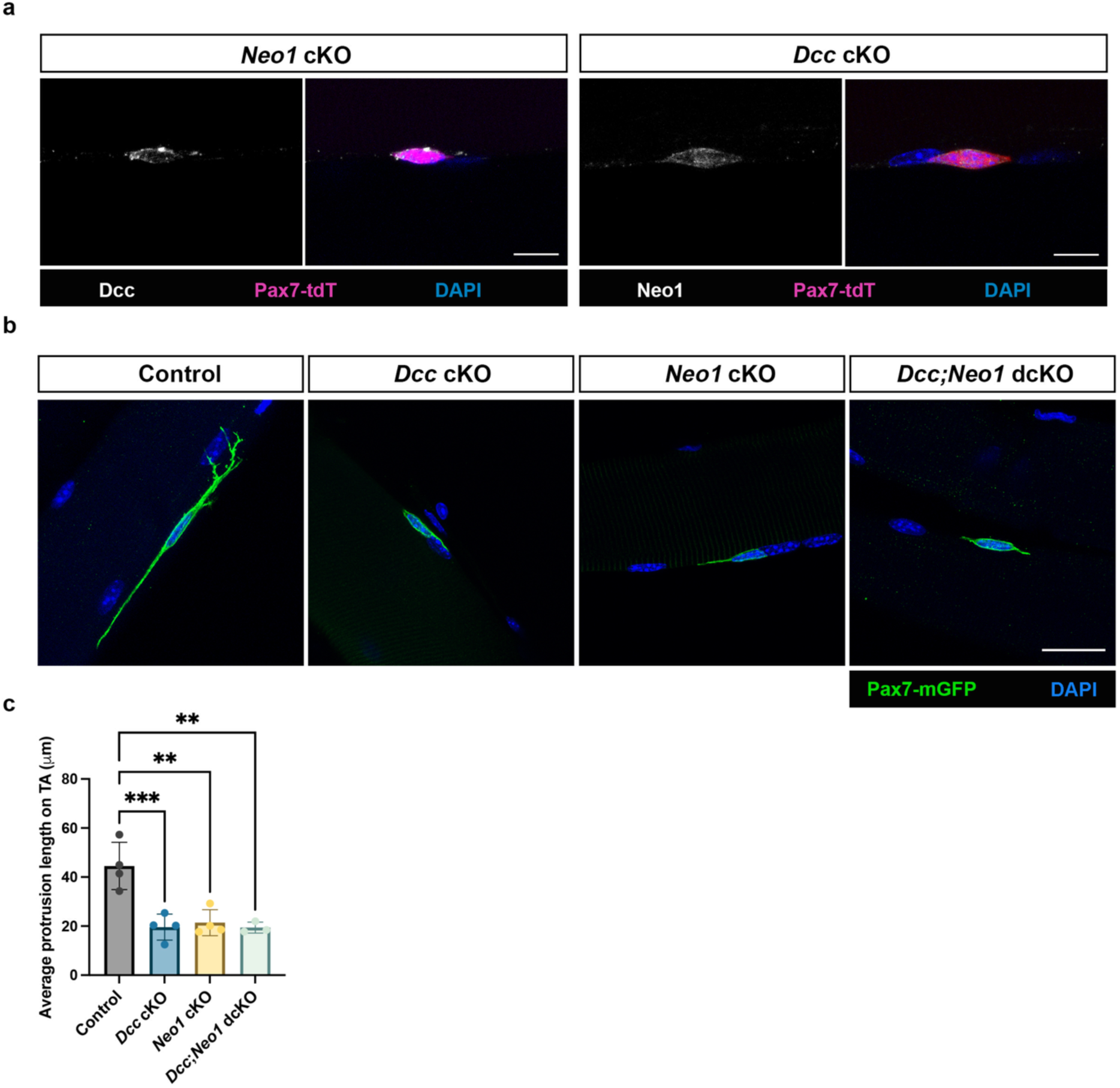
Short MuSC protrusions in the TA muscles of *Dcc* cKO, *Neo1* cKO, and *Dcc;Neo1* dcKO mice. **a,** Expression of Dcc protein in *Neo1* cKO MuSCs and expression of Neo1 protein in *Dcc* cKO MuSCS. Single myofibers from the indicated mutant line were immunostained with antibodies to tdTomato to reveal MuSCs, and Dcc or Neo1 as indicated. Scale bar, 10μm. **b,** Immunofluorescence analysis of protrusion length in MuSCs from control, *Dcc* cKO, *Neo1* cKO, and *Dcc;Neo1* dcKO *Pax7^mG^*mice. TA muscles were dissected and immediately fixed and immunostained with antibodies to GFP. Scale bar, 20μm. **c,** Quantification of average MuSC protrusion length in TA muscles represented in (**b**). Each point represents the average protrusion length of at least 50 MuSCs from an individual mouse.

**Extended Data Fig. 3.**
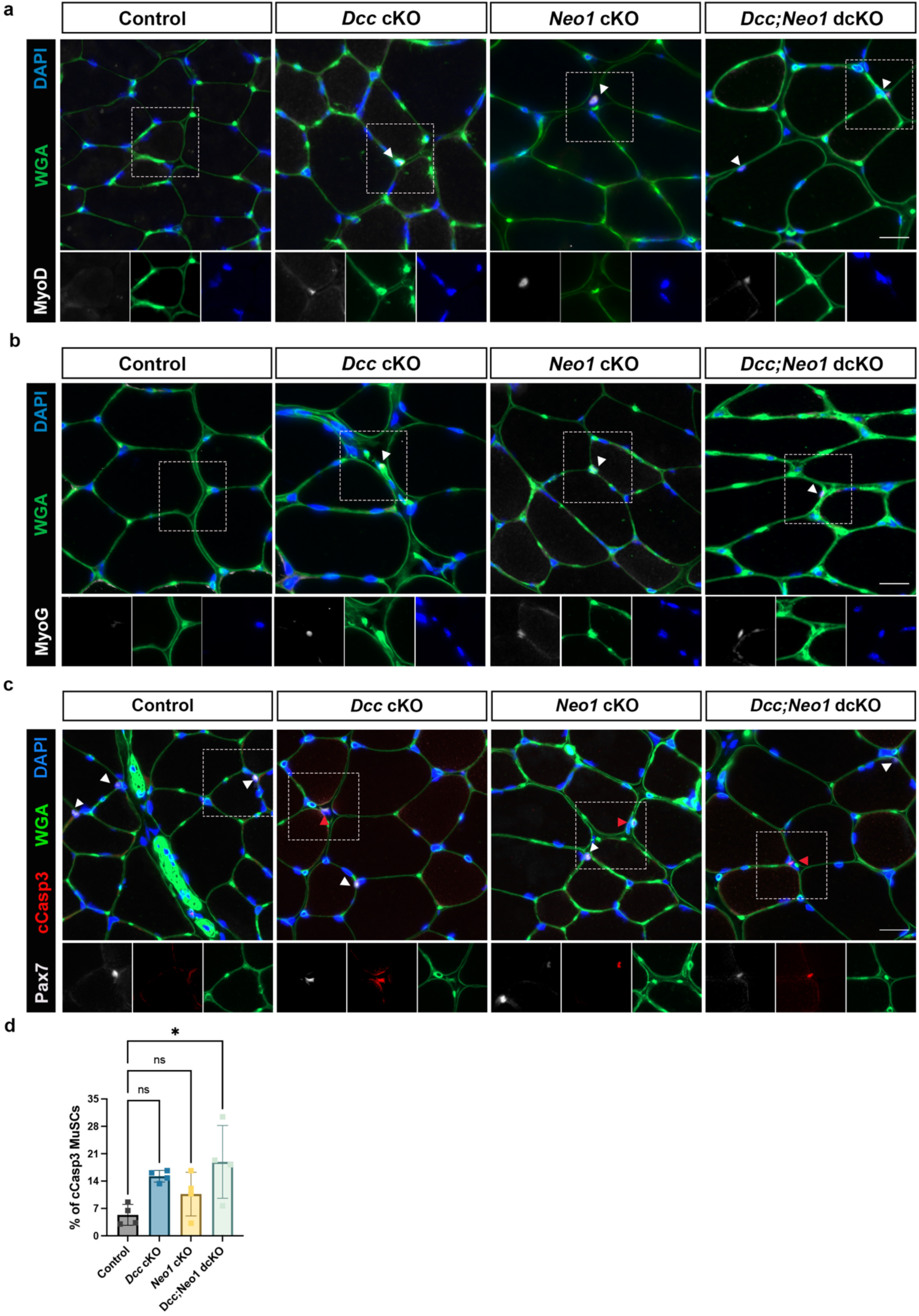
Production of MyoD, MyoG, and cleaved Caspase-3 by MuSCs in *Dcc* cKO, *Neo1* cKO, and *Dcc;Neo1* dcKO mice. **a-c,** Immunofluorescence analysis of TA muscle sections from control, *Dcc* cKO, *Neo1* cKO, and *Dcc;Neo1* dcKO mice. Sections were stained with antibodies to MyoD, myogenin (MyoG), and Pax7 plus cleaved Caspase-3 (cCasp3) as indicated. WGA staining outlines myofibers. Scale bars, 20μm. **d,** Quantification of the percentage of cCasp3^+^ Pax7^+^ MuSCs in sections represented in (c). White arrowheads represent Pax7^+^ MuSCs, and red arrowheads represent cCasp3^+^/Pax7^+^ cells.

## Supplementary Videos

**Supplementary Video 1.** Video of a single tdTomato-labeled MuSC retracting a cellular protrusion. Still images from the video are displayed in Fig. 1b.

**Supplementary Video 2.** Video of a single tdTomato-labeled MuSC retracting and then extending cellular protrusions. Still images from the video are displayed in Fig. 1c.

**Supplementary Video 3.** Video of a single tdTomato;EB1-EGFP-labeled MuSC demonstrating a dynamic filopodium. Still images from the video are displayed in Fig. 1e.

**Supplementary Video 4.** Video of a tdTomato-labeled MuSC (upper cell) demonstrating dynamic filopodia.

**Supplementary Video 5.** Video of a single tdTomato-labeled MuSC extending cellular protrusions in response to Netrin-1. Still images from the video are displayed in Fig. 3a.

**Supplementary Video 6.** Video of a single tdTomato-labeled MuSC failing to extend cellular protrusions in response to Netrin-1 when co-treated with the Rac inhibitor NSC23766. Still images from the video are displayed in Fig. 3a.

**Supplementary Video 7.** Video of a single tdTomato-labeled MuSC failing to extend cellular protrusions in response to Netrin-1 when co-treated with the Arp2/3 inhibitor CK-666. Still images from the video are displayed in Fig. 3a.

**Supplementary Video 8.** Video of a single tdTomato-labeled *Dcc* cKO MuSC failing to extend cellular protrusions in response to Netrin-1. Still images from the video are displayed in Fig. 5a.

**Supplementary Video 9.** Video of a single tdTomato-labeled *Neo1* cKO MuSC failing to extend cellular protrusions in response to Netrin-1. Still images from the video are displayed in Fig. 5b.

## Notes

### Competing Interest Statement

The authors have declared no competing interest.

### Summary of Updates

Additional results and text have been added.

